# Genetic recombination of poliovirus facilitates subversion of host barriers to infection

**DOI:** 10.1101/273060

**Authors:** Ashley Acevedo, Andrew Woodman, Jamie J. Arnold, Ming Te Yeh, David Evans, Craig E. Cameron, Raul Andino

**Affiliations:** Department of Microbiology and Immunology, University of California, San Francisco 94122-2280, USA; Biomedical Sciences Research Complex, University of St. Andrews, UKKY16 9ST; Department of Biochemistry and MolecularBiology, The Pennsylvania State University, University Park, PA 16802 USA

**Author notes:** **Corresponding Author:** Correspondence should be addressed to R.A.

## Abstract

The contribution of RNA recombination to viral fitness and pathogenesis is poorly defined. Here, we isolate a recombination-deficient, poliovirus variant and find that, while recombination is detrimental to virus replication in tissue culture, recombination is important for pathogenesis in infected animals. Notably, recombination-defective virus exhibits severe attenuation following intravenous inoculation that is associated with a significant reduction in population size during intra-host spread. Because the impact of high mutational loads manifests most strongly at small population sizes, our data suggest that the repair of mutagenized genomes is an essential function of recombination and that this function may drive the long-term maintenance of recombination in viral species despite its associated fitness costs.

**Significance Statement:** RNA recombination is a widespread but poorly understood feature of RNA virus replication. For poliovirus, recombination is involved in the emergence of neurovirulent circulating vaccine-derived poliovirus, which has hampered global poliovirus eradication efforts. This emergence illustrates the power of recombination to drive major adaptive change; however, it remains unclear if these adaptive events represent the primary role of recombination in virus survival. Here, we identify a viral mutant with a reduced rate of recombination and find that recombination also plays a central role in the spread of virus within animal hosts. These results highlight a novel approach for improving the safety of live attenuated vaccines and further our understanding of the role of recombination in virus pathogenesis and evolution.

## Introduction

Recombination in RNA viruses enables exchange of genetic material through a copy-choice mechanism in which the viral polymerase switches templates during replication (1). This process is poorly defined mechanistically but may involve local sequence complementarity (1, 2) or the biphasic selection of genomes with increased fitness (3), resulting in the formation of recombinants lacking insertions and deletions. RNA recombination appears to be a general phenomenon in RNA viruses, though its frequency varies considerably (reviewed in 4). In poliovirus, the first RNA virus in which genetic recombination was observed (5, 6), the frequency of recombination across the genome is estimated to be as high as 10-20% (1, 7, 8).

Despite the prevalence of recombination among RNA viruses, its biological role is poorly understood. RNA recombination is thought to benefit viruses by accelerating combination of beneficial mutations from different lineages (9, 10) and/or by purging of deleterious mutations (11, 12). Studies in HIV-1 have shown that recombination contributes to the acquisition of multidrug resistance (13, 14) and immune escape (15, 16), demonstrating the potential advantage of recombination in speeding up production of adaptive genotypes. However, because RNA viruses exhibit high mutation rates and thus may readily generate advantageous combinations by mutation alone, the relative contribution of recombination is unclear. Alternatively, high mutational loads in RNA viruses may necessitate purging of deleterious mutations to maintain population fitness. This is particularly important in small populations subject to high levels of drift where populations can undergo irreversible accumulation of deleterious mutations in a process called Muller’s Ratchet (11, 17, 18). Though not shown for recombination directly, a study of the RNA bacteriophage Phi6 demonstrated that reassortment, an analogous method of genetic exchange in segmented RNA viruses, slows the decline in fitness characteristic of Muller’s Ratchet (19). While experimental evidence supports both proposed roles of recombination, the direct impact of recombination on viral fitness and pathogenesis and the mechanisms that confer those effects remain unclear.

In this study, we isolated a poliovirus variant defective for recombination and use this variant to examine the effects of recombination in RNA viruses. We found that in tissue culture, where replication efficiency is paramount, recombination imposes a fitness cost. However, in a mouse model for poliovirus pathogenesis, the effect of recombination is dependent on the route of infection. This dependency is associated with the severity of host bottlenecks to virus entry into the central nervous system. Strikingly, we found that recombination defective virus inoculated intravenously, the route of infection that imposes the most stringent host bottleneck, exhibits severe attenuation of viral virulence, suggesting that recombination is a critical mechanism for overcoming host bottlenecks. It is thus possible that the reduction of population size during repeated bottleneck events may require effective RNA recombination to ensure viral survival. These findings imply that ecological forces within the infected individual underlie the advantage of recombination and may ultimately drive its evolutionary maintenance in RNA viruses despite its replicative cost.

## Results

### Isolation of a recombination deficient poliovirus

To identify determinants modulating the recombination rate of poliovirus we designed a genetic system composed of a replication-competent, recombinant poliovirus that encodes enhanced green fluorescent protein (eGFP) within the viral polyprotein. To release eGFP from the polyprotein and ensure correct proteolytic processing of the endogenous viral proteins, eGFP is flanked by proteolytic cleavage sites recognized by the poliovirus 2A protease (Fig. 1A). While the eGFP-expressing virus is able to produce all of its constituent proteins and proceed with replication normally, previous work has shown that this virus is genetically unstable, where homologous recombination at nucleotide sequences encoding the 2A cleavage sites results in precise deletion of the inserted sequence (20). We reasoned that genetic variants reducing the rate of homologous recombination would increase the stability of eGFP-expressing virus.

**Figure 1.**
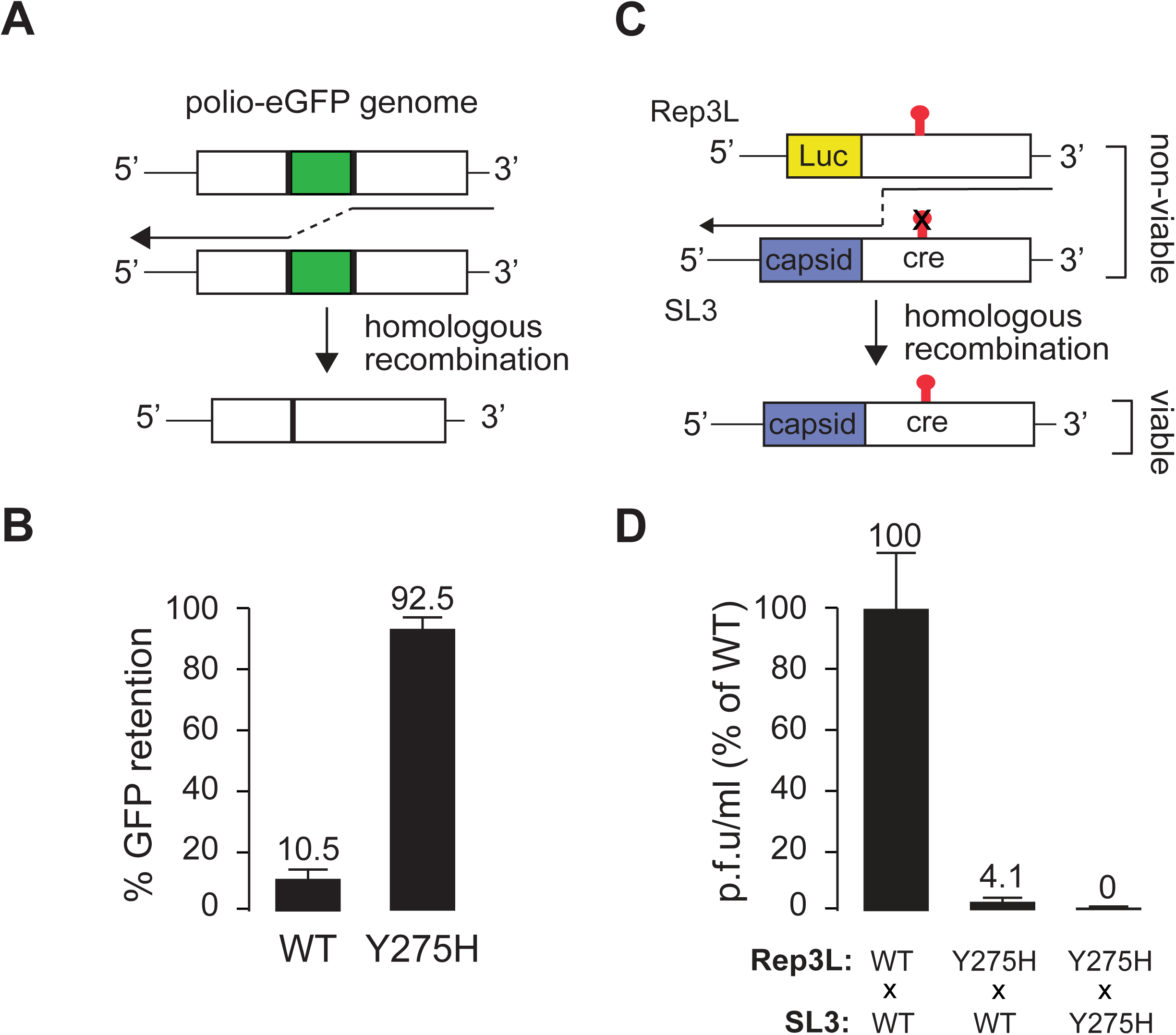
A hybrid screen:selection identifies a recombination deficient virus containing the mutation Y275H in 3D^pol^. (A) Hybrid screen:selection scheme. Recombination at duplicate 2A cleavage sites (black, GATCTGACCACATATGGATTCGGACAC) can lead to excision of eGFP (green) from the genome. Inability to recombine leads to high rates of GFP retention. (B) Percent eGFP retention by wild type and 3D^pol^ Y275H virus is measured by limiting dilution of virus produced by electroporation of infectious RNA. Error bars represent 95% binomial confidence intervals. (C) Schematic representation of CRE-REP assay. SL3 and Rep3L RNAs are co-transfected into mouse cells. SL3 and Rep3L are non-viable poliovirus RNAs with defects in an essential replication element (CRE) or that lack structural proteins, respectively. Progeny generated through recombination between the capsid-coding region and CRE are viable, containing both a functional CRE and structural genes. (D) Relative titers of viable, recombinant progeny after a single round of replication. Titers are normalized to Rep3L and SL3 RNAs containing wild type viral polymerase. Error bars represent standard deviations of three replicates.

To identify recombination-deficient poliovirus variants, eGFP chimeric viruses were cloned by limiting dilution and screened for eGFP expression. Clones expressing eGFP were isolated and further propagated. This cycle was repeated until we isolated a viral strain that stably expressed eGFP. This strain contained two substitutions: isoleucine 37 to valine (I37V) within the 2C protein and tyrosine 275 to histidine (Y275H) in the viral RNA-dependent RNA polymerase (3D^pol^). Each substitution was individually engineered into the initial chimeric eGFP virus. While I37V conferred a modest (2.7 fold) increase in eGFP retention compared to wild type (not shown), Y275H increased eGFP retention by 8.8 fold compared to wild type, greater than 90% retention per replication cycle (Fig. 1B). Previously, this process was used to identify the aspartic acid 79 to histidine (D79H) substitution in 3D^pol^, which exhibits an intermediate eGFP retention rate of 56.5%, a 5.4 fold increase over wild type (21, 22). Given these results and previous observations suggesting that recombination requires RNA synthesis (1) and that purified poliovirus 3D^pol^ is able to mediate effective template switching *in vitro* (23), we selected the Y275H variant for further studies.

### Y275H exhibits a bona fide recombination defect

To further examine the effect of Y275H on recombination, we employed a cell culture-based recombination assay (3). This assay utilizes two parental viral RNAs that are independently unable to generate viable progeny: SL3, containing a mutated *cis*-acting replication element (CRE) that prevents positive strand synthesis (24) and Rep3L, a replicon that does not encode structural proteins. Following co-transfection of SL3 and Rep3L *in vitro* transcribed RNA, viable progeny are produced if recombination between defective RNAs takes place at any site between the structural proteins and defective CRE (3) (Fig. 1C). Strikingly, introduction of the Y275H mutation into either the 3D^pol^ of Rep3L alone or into both Rep3L and SL3 dramatically reduces the number of viable recombinant progeny (Fig. 1D). In comparison, introduction of the D79H mutation yields an intermediate number of viable recombinant progeny (Fig. S1), consistent with its lower rate of eGFP retention.

We also examined the capacity of purified Y275H 3D^pol^ to mediate recombination in a reconstituted system using a template-switching assay (25). In this assay, elongation complexes composed of 3D^pol^ and a symmetrical, heteropolymeric primer-template substrate (sym/sub-U) (26) are extended by the addition of nucleotides in the presence of an excess of an RNA acceptor template that is partially homologous to the sym/sub-U template strand (Fig. 2A). In reactions containing wild type polymerase, the elongating polymerase switches from the sym/sub-U template to the RNA acceptor template generating a longer recombinant transfer product (Fig. 2B). These products are not observed in the absence of RNA acceptor template or in the presence of RNA acceptor that is non-complementary to the extended sym/sub-U template (strong stop product, Figs. 2A and S2). In contrast, under conditions in which Y275H 3D^pol^ produces comparable amounts of strong-stop product as wild-type enzyme, the Y275H polymerase exhibits a dramatic reduction in the appearance of the longer product of recombination (Fig. 2B). These results demonstrate that Y275H has a defect in its ability to switch templates that is unrelated to its elongation function.

**Figure 2.**
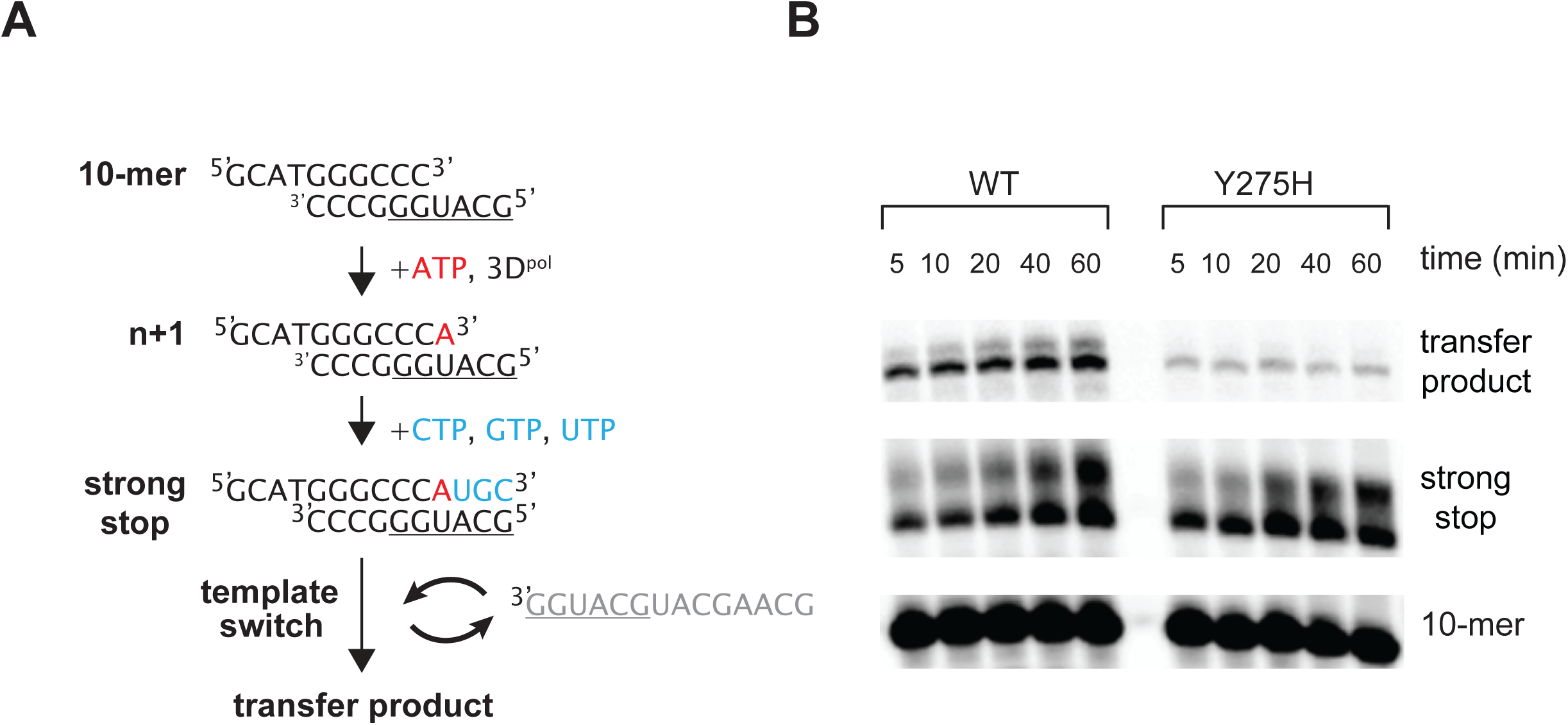
A reconstituted assay reveals the template switching defect of Y275H. (A) Scheme for *in vitro* template-switching assay. ATP and 3D^pol^ are combined with sym/sub-U (black) to produce 3Dpol-sym/sub-U elongation complexes. Elongated complexes, which are partially complementary to the RNA acceptor template (grey), can switch templates to produce longer nascent strands (transfer product). In addition to the transfer product shown, other junctions can also be observed after cloning and sequencing (JJA and CEC, unpublished). (B) 500 µM CTP, GTP and UTP and 60 µM RNA acceptor are added to 3Dpol-sym/sub-U elongation complexes formed in the presence of 500 µM ATP. Reactions were quenched after 5, 10, 20, 40 and 60 minutes. Transfer product results from template switch with RNA acceptor.

Furthermore, because the fidelity of 3D^pol^ potentially associates with the rate of recombination (3, 25, 27), we examined the Y275H mutant virus for changes in replication fidelity. Previous work has shown 3D^pol^ misincorporates the RNA mutagen ribavirin (28), where the fidelity of 3D^pol^ determines the efficiency of ribavirin utilization. The high fidelity G64S 3D^pol^ is less efficient than wild type at incorporating ribavirin and thus is resistant to its mutagenic effects (29), whereas the low fidelity H273R 3D^pol^ incorporates ribavirin more efficiently and thus is highly sensitive to ribavirin’s mutagenic effects (30). We grew wild type, Y275H, G64S and H273R mutant viruses in increasing concentrations of ribavirin (Fig. S3). In this assay, the Y275H virus exhibits ribavirin sensitivity similar to that of the wild type virus, indicating that Y275H does not significantly alter replication fidelity. Thus, while in some cases fidelity and recombination rate may be linked, they can also be modulated independently, as in the case of the Y275H mutant.

### Recombination imposes a fitness cost in cell culture

We examined the growth characteristics of Y275H by measuring its replication kinetics over a single replication cycle in human and mouse cell lines (HeLa S3 and L20B, human poliovirus receptor-expressing murine cells; 31). Though the virus displays a slight lag in replication relative to the wild type midway through infection (4-6 hours post infection, Figs. 3A and B), titers reach similar levels by the end of the infection cycle (8 hours post infection). Because this assay is relatively insensitive to fitness differences between strains, we performed competitions between Y275H and the wild type virus. We infected monolayers of HeLa S3 or L20B cells with a 1:1 mixture of wild type and Y275H virus at a low multiplicity of infection (m.o.i.) and allowed the infection to proceed for a single replication cycle (8 hours). The low m.o.i. ensures that only a single infectious virus enters one cell and thus, that replication of the virus is not affected by complementation. Progeny collected at the end of the infection cycle was then used to initiate another round of low m.o.i. infection. We repeated this procedure 4 times and determined the proportion of each virus type at each passage by digital droplet reverse transcription PCR using TaqMan probes specific for either wild type or Y275H virus. In both human and murine cells, Y275H rapidly outcompetes wild type (Figs. 3C and D), increasing its relative abundance at each passage. These results are remarkably consistent with direct measurement of relative fitness using CirSeq, a population sequencing-based approach, which showed that Y275H has a 26% (± 2%) fitness advantage relative to wild type (32). The D79H mutation also shows a fitness advantage relative to wild type, though that advantage is weaker than that observed for Y275H (Fig. S4). Thus, regardless of host cell type, viral recombination has a cost for poliovirus replication, resulting in a significant reproductive disadvantage.

**Figure 3.**
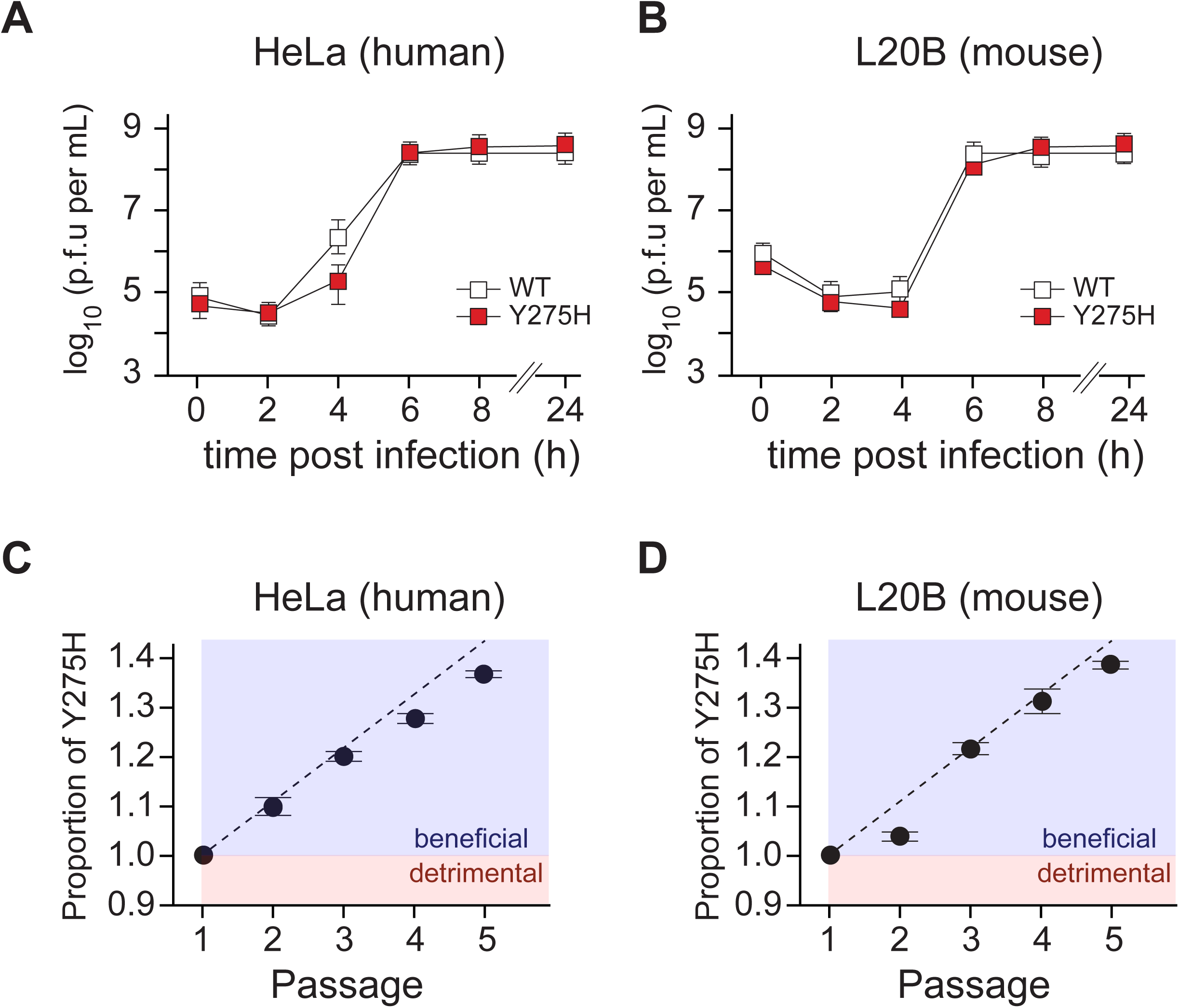
Recombination reduces fitness in cultured cells. (A and B) Growth kinetics of wild type (white) and Y275H (red) virus in (A) HeLa S3 and (B) L20B cells. Error bars represent standard deviations of three replicates. (C and D) Competition of wild type and Y275H virus in (C) HeLa S3 and (D) L20B cells. Viruses were mixed 1:1 and serially passaged at low m.o.i. Proportions of each virus type were determined by TaqMan RT-PCR in digital droplets. Dashed line represents a model of growth of Y275H relative to wild type based on the 26% fitness advantage observed in Acevedo et al., 2014. Error bars represent standard deviations of three replicates.

### Recombination alters pathogenesis in an infection route-dependent manner

Previous efforts to examine the impact of recombination on virus pathogenesis using the D79H mutation revealed a minor reduction of poliovirus virulence (21). Given the significantly stronger deficiency in recombination exhibited by the Y275H mutation, we sought to gain a clearer understanding of the role of recombination in virus pathogenesis. To that end, we infected poliovirus susceptible mice (33) intracranially, intramuscularly and intravenously with either wild type or Y275H virus. Infection of these mice leads to acute central nervous system (CNS) disease marked by paralysis and, in the majority of cases, eventual death. For direct infection of the CNS via the intracranial route, we observed no difference in time-to-death (p = 0.287, log-rank test) or survival (p = 0.586 at 15 days post infection (d.p.i.), fisher’s exact test) between wild type and Y275H (Fig. 4A). For intramuscular infection, both wild type and Y275H are similarly virulent (p = 1 at 14 d.p.i, fisher’s exact test), causing invasion of the CNS and fatal paralytic disease in approximately the same number of mice, however, Y275H displays a statistically significant increase in time-to-death (p = 0.0258, log-rank test) (Fig. 4B). Strikingly, intravenous infection results in a significant attenuation of virulence (p = 7.0x10^-4^, fisher’s exact test) and consequently a significant increase in time-to-death (p = 2.3x10^-6^, log-rank test) for Y275H relative to wild type virus (Fig. 4C). Thus, despite its replicative advantage in cultured cells, Y275H is less efficient at invading the CNS than wild type virus. This observation suggests that recombination in wild type poliovirus confers a significant advantage during virus spread within the infected individual, which compensates for its replicative disadvantage.

**Figure 4.**
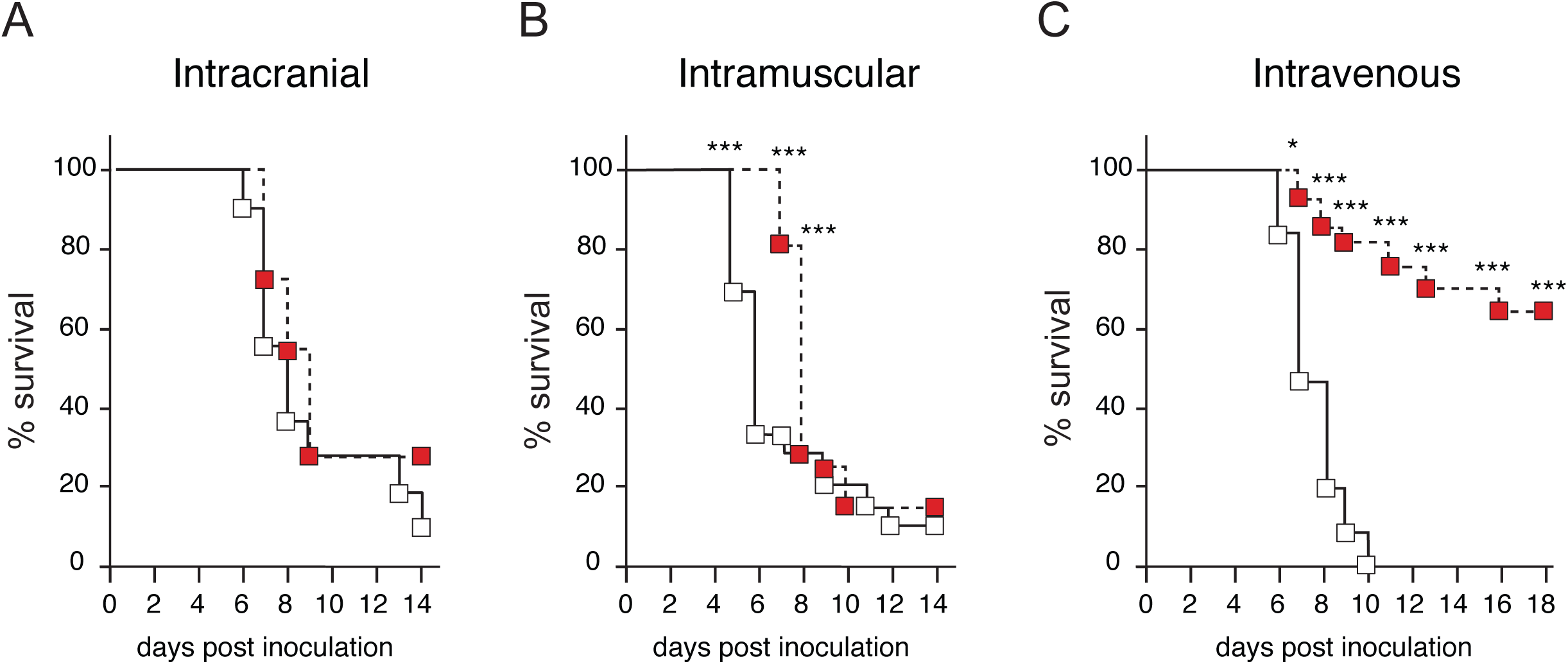
Recombination alters pathogenesis in an infection route dependent manner. (A) Survival of mice infected intracranially. Eleven 2 week old PVR transgenic mice were injected with 5x10^4^ plaque forming units (p.f.u.) of wild type (white) or Y275H (grey) virus. We observe no difference in time-to-death (p = 0.287, log-rank test). (B) Survival of mice infected intramuscularly. Twenty-five 8-10 week old PVR transgenic mice were injected with 10^6^ plaque forming units (p.f.u.) of wild type (white) or Y275H (grey) virus. The difference in time-to-death is significant (p = 0.0258, log-rank test). (C) Survival of mice infected intravenously. Fifteen 6 week old mice were injected with 3x10^8^ p.f.u. of wild type (white) or Y275H (grey) virus. The difference in time-to-death is significant (p = 1.02x10^-6^, log-rank test). Virulence is significantly reduced (p = 7.0x10-4 at 18 days post infection, fisher’s exact test). (A-C) Time points with significant differences in survival by fisher’s exact test are noted as follows: * = p < 0.05, ** = p < 0.01 and *** = p < 0.001.

The genetic advantage conferred by recombination is highly dependent on the route of infection, where there is no advantage upon intracranially infection but a highly significant advantage upon intravenous infection. A major distinction between these infection routes is the number of different tissues the virus must transit before it can access the neuronal tissues where it replicates and causes clinical symptoms. Intracranial infection delivers poliovirus directly to the CNS, whereas poliovirus injected intramuscularly must first enter peripheral nerves at neuromuscular junctions near the injection site and then access the CNS via retrograde axonal transport (34–36). Further, intravenous inoculation first induces a transient viremia leading to virus invasion of non-neuronal tissues, including skeletal muscle, from which, like intramuscular infection, it gains access to neuromuscular junctions and subsequently the CNS (34). The additional steps required to access the CNS via the intramuscular and intravenous routes introduce host barriers to infection through a variety of mechanisms (37), which result in population bottlenecks. As such, studies examining population bottlenecks in poliovirus infected mice have shown that, while there is virtually no population bottleneck imposed by intracranial infection, bottlenecks for intramuscular and intravenous infection are significant with intravenous infection experiencing the most severe host bottleneck of the routes employed here (38, 39). The tight association between the stringency of population bottlenecks and the advantage of recombination suggests that recombination provides a critical mechanism for overcoming population bottlenecks caused by host barriers to infection.

## Discussion

Here we isolate a viral variant shown to modify the rate of RNA recombination. This variant affects neither the ability of 3D^pol^ to elongate nor its fidelity, highlighting that recombination rates can be selected independently of other essential functions of the viral polymerase. Our results suggest that recombination has no apparent benefit in cell culture, in fact, it has a significant fitness cost, and thus, it is in principle unclear why recombination is so prevalent among positive-stranded RNA viruses. Strikingly, recombination is necessary for entry into the CNS and, consequently, recombination defective poliovirus is attenuated in an animal model of infection. Thus recombination provides a fitness advantage in the complex environment of the infected individual, specifically by enabling virus to overcome host barriers to infection. These results stand in contrast to assertions that recombination is merely a by-product of evolution acting on other functions of the polymerase or other aspects of virus biology (4) and indicate that recombination provides an evolutionary advantage to virus survival in nature.

The recombination-defective variant we describe, Y275H, is located at a position in the structure that permits the histidine substitution to modulate template interactions (Fig. 5). When one considers the physical nature of template switching, two possibilities have been put forward. One possibility is that the polymerase interacts with the nascent RNA much more than with the template and the polymerase-nascent RNA complex moves from one template to another (1, 23). Another possibility is that a homologous acceptor template adds to form a three-stranded intermediate (44, 45). In either case, interactions of the polymerase with the template would need to be enhanced. If the first mechanism is operative biologically, then enhanced interactions with template might be expected to increase processivity, the average number of nucleotides incorporated into nascent RNA during a single binding event.

**Figure 5.**
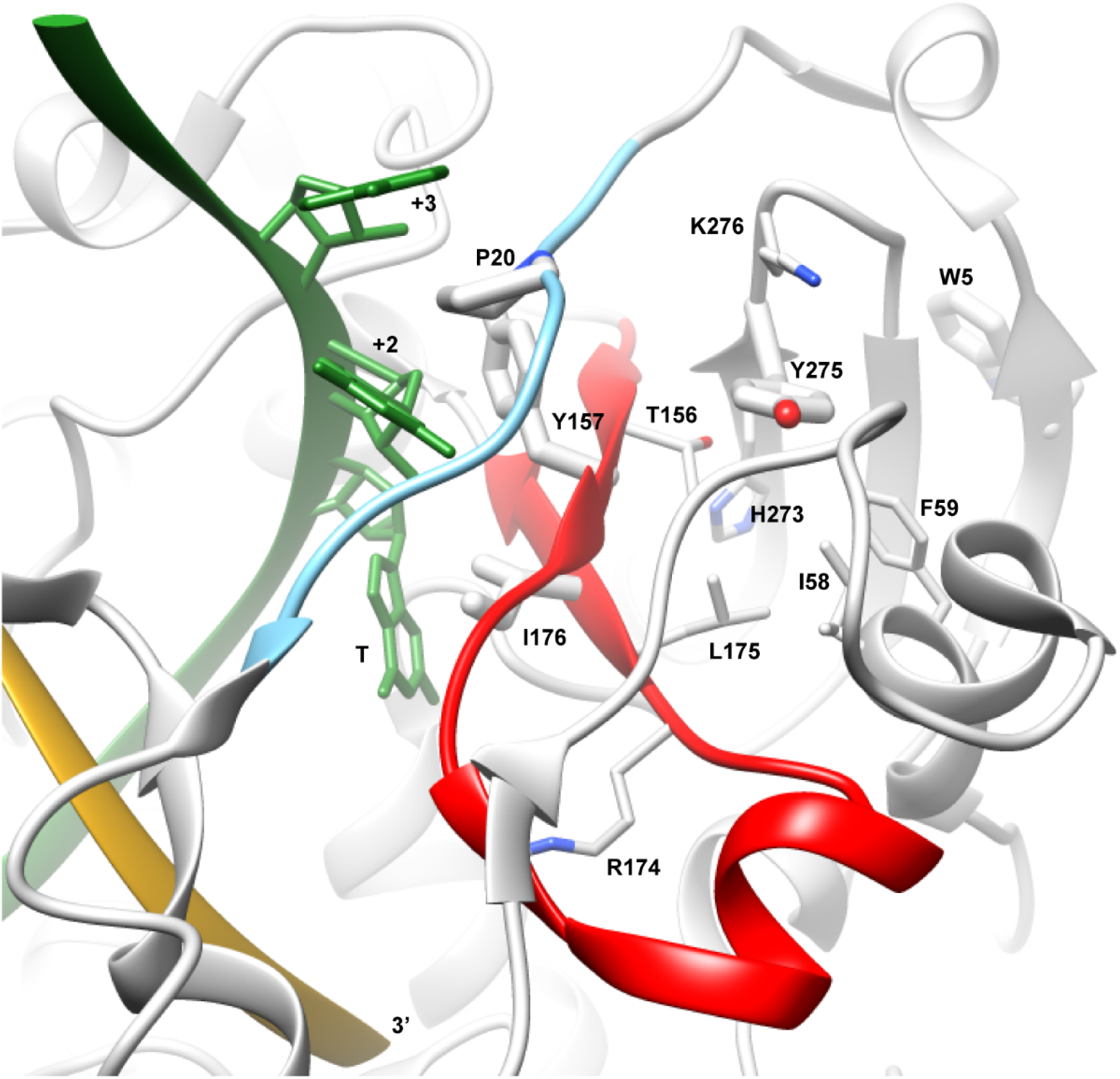
How might changes to Y275 impact interactions with an elongation complex? Shown is the environment surrounding Y275 in PV RdRp-RNA binary complex (PDB 3OL6; 40). Previous studies have shown that residues 18-26 (cyan) are dynamic in the absence of primer-template (41). Movement of this element is required for template binding (42). In fact, P20 intercalates between the +2 and +3 nucleotides of template RNA. Clearly, the disposition of the 18-26 element will be influenced by conserved structural motif F (red). Motif F also contributes to template binding. Y157 interacts with the +2 template position; I176 stacks against the templating base (denoted T). Y275 is located at the end of a strand of a β-sheet and is buried in a pocket comprised of residues I58, F59, I175 and the aliphatic portion of K276. The position of the β-sheet will influence the position of motif F, which, in turn will influence the 18-26 element and interactions with template. Interestingly, substitutions at W5, another component of the β-sheet, are known to modulate the stability of the RdRp on a primed template (43). It is therefore possible that the H275 derivative perturbs the interaction with template RNA.

Enhanced processivity might also explain the observation that Y275H is more fit than wild-type virus in cell culture. Movement of a polymerase-nascent RNA complex from one template to another is unlikely to occur with 100% efficiency. Therefore, it is likely that template switching will yield some quantity of prematurely terminated (anti)genomes. Enhanced processivity will yield fewer prematurely terminated genomes--that is, more viable progeny.

Virus replication in animals introduces additional challenges to infection. In this more complex environment, we show that, depending on the route of infection, recombination can enhance viral fitness, facilitating invasion of the CNS and pathogenesis. Specifically, we find no detectable advantage of recombination upon intracranial infection, a moderate advantage upon intramuscular infection, where recombination defective virus exhibits a significant lag in pathogenesis, and a strong advantage upon intravenous infection, resulting in severe attenuation of the recombination-defective virus. These routes of infection tightly associate with the stringency of population bottlenecks caused by host barriers to infection (38, 39), where recombination is most advantageous at smaller effective population sizes. In these small populations, it is hypothesized that fit genotypes are highly vulnerable to loss through random sampling and mutagenesis. Repeated loss of these fit genotypes leads to a process termed Muller’s ratchet in which populations undergo deterioration of fitness from the accumulation of deleterious mutations (11, 17, 18). Recombination reverses this process by restoring genetically undamaged, fit genotypes, in effect, purging deleterious mutations from the population (11, 17, 19). Our finding that virus recombination is more advantageous in smaller populations is consistent with reversal of the effects of Muller’s ratchet, suggesting that the repair of mutagenized genomes is a critical process for viruses in overcoming barriers to infection and mediating pathogenesis. Importantly, in the context of this animal model for poliovirus infection, the ability to effectively transit host bottlenecks results in enhanced virulence. While it is unclear if enhanced virulence is adaptive (reviewed in 46), these results offer a paradigm to explain the genetic mechanisms that enable viruses overcome bottlenecks.

Viruses routinely encounter population bottlenecks, a result of intra-host barriers (e.g. innate and adaptive immune responses or physical barriers like the blood-brain barrier) and host-to-host transmission (e.g. respiratory droplets), as inevitable steps in their life cycles. Further, the high mutation rates characteristic of RNA viruses likely exacerbate the pressures of survival in these small populations (47), threatening population extinction by lethal mutagenesis. These challenges and the capacity for recombination to alleviate them provide an explanation of why recombination in poliovirus is conserved, despite its replicative disadvantage. It also likely explains the prevalence of recombination across a wide range of viral species (reviewed in 4). Though our findings indicate that the primary advantage of genetic recombination in acute infection is in its capacity to preserve genetic information and fitness, it is likely that over longer evolutionary time scales recombination also accelerates the creation of new combinations of alleles and phenotypic traits, enabling adaptation to novel environments (e.g. new tissues or hosts, changes in immune responses, therapeutic interventions). Moreover, manipulation of this fundamental process may help to understand the principles modulating pathogenesis and provide novel avenues to target and attenuate viral pathogens. Finally, the identification of recombination determinants within 3D^pol^ is central to the copy-choice recombination mechanism and may open novel opportunities to further explore the mechanism of this important, but poorly understood process.

## Materials and Methods

### Plasmids and *in vitro* transcription

prib(+)XpA (48) contains the full-length poliovirus type 1 Mahoney cDNA. Y275H, G64S (49, and H273R (30) were cloned by quickchange mutagenesis using single primers (Y275H: 5’- AACCACTCACACCACCTGCACAAGAATAAAACATACTGT-3’; G64S: 5’- ATTTTCTCCAAGTACGTGTCAAACAAAATTACTGAAGTG-3’; H273R: 5’- TACCTAAACCACTCACACAGACTGTACAAGAATAAAACA-3’) from IDT. eGFP flanked by nonhomologous 2A cleavage sites was cloned from pMov2.8-EGFP (33) into the prib(+)XpA backbone. The 5’ 2A cleavage site was altered by overlap extension PCR using primers (external-2439 F: 5’-TGCGAGATACCACACATATAGAGC-3’; external-4393 R: 5’-AGGGGCAAACCTCTTAGACTGGATGGATAAC-3’; internal-2A cleavage site F: 5’- GATCTGACCACATCTGGATTCGGACACGGCGGAGGTGGGGGAGGTGAATTC-3’; internal-2A cleavage site R: 5’- GTGTCCGAATCCATATGTGGTCAGATCCTTGGTGGAGAGGGGTGTAAGCGT-3’) from IDT. The resulting plasmid, prib(+)XpA Polio-eGFP, contains eGFP flanked by homologous 2A cleavage sites. pRLucWT and pT7Rep3L are sub-genomic replicons of poliovirus type 1 Mahoney and type 3 Leon, repectively, containing a luciferase reporter gene in place of structural proteins, as previously described (51, 3). pT7/SL3 contains full-length poliovirus type 3 Leon cDNA with 8 synonymous mutations in the *cis*-acting replication element (CRE), as previously described (24).

To produce RNA, prib(+)XpA based constructs were linearized with MluI and *in vitro* transcribed under the following conditions: 400 mM HEPES pH 7.5, 120 mM MgCl_2_, 10 mM Spermidine, 200 µM DTT, 7.5 mM each of ATP, CTP, GTP and UTP, 1 µg linearized DNA and 1 µl purified T7 polymerase in 25 µl total volume. pT7Rep3L and pT7/SL3 were linearized with SalI and pRLucWT was linearized with ApaI and then *in vitro* transcribed using T7 Polymerase (Fermentas) following the manufacturer’s protocol.

### Cells and viruses

HeLa S3 (ATCC, CCL2.2) and L20B cells were propagated in DMEM High Glucose/F12 medium supplemented with 10% newborn calf serum (SIGMA) and 1x Pen Strep Glutamine (Gibco) at 37°C. L929 (murine) and HeLa cells were propagated in DMEM with 10% heat inactivated fetal calf serum and 1x Pen Strep.

Wild type, Y275H, G64S, H273R and eGFP containing poliovirus were generated by electroporation of HeLa S3 cells with *in vitro* transcribed RNA with a BTX electroporator using the following settings: 300 V, 1000 µF, 24 Ω in a 0.4 cm electroporation cuvette (BTX). Cells were incubated 16 hours at 37°C then frozen and thawed 3 times and cleared at 3500 rpm for 5 minutes to produce initial viral stocks.

Wild type and Y275H virus for animal infections were produced by passaging initial viral stocks 3 times at a multiplicity of infection (m.o.i.) 1 until total cytopathic effect (CPE). The third passage was performed in medium lacking serum.

### Selection of recombination deficient virus

An initial viral stock of Polio-eGFP was titered in HeLa S3 by TCID_50_ and then plated in 96-well plates at .25 TCID_50_ per well. Plates were scanned on a TECAN Safire plate reader using the following settings: measurement mode - fluorescence bottom, excitation wavelength - 488 nm, emission wavelength - 509 nm, excitation bandwidth - 2.5 nm, emission bandwidth - 2.5 nm, gain - 100. Supernatant from wells positive for eGFP was collected, combined and then titered by TCID_50_. This cycle was repeated until the ratio of eGFP positive:CPE positive wells was greater than 0.9. Variants fixed in this population were recloned into the Polio-eGFP vector, *in vitro* transcribed and then electroporated into HeLa S3 cells. To measure the rate of retention of eGFP, virus from these cells was diluted into 96-well plates at .25 TCID_50_ per well and monitored fluorescence and CPE.

### One-step growth assay

HeLa S3 cells were infected in triplicate with wild type or Y275H at m.o.i. 5 for 30 minutes, washed 2 times with PBS and covered with growth medium. Cells were frozen at 0, 2, 4, 6, 8 and 24 hours post infection. Cells were frozen and thawed 3 times and cleared for 5 minutes at 21000 g. Supernatants were titered by plaque assay.

### Cell culture competition assay

For digital droplet-based detection, HeLa S3 and L20B cells were coinfected with wild type and Y275H virus at m.o.i. 0.05 each for 8 hours in triplicate. Cells were frozen and thawed 3 times and cleared at 3500 rpm for 5 minutes. The viral supernatants were titered by plaque assay and passaged further at low m.o.i. for 8 hours. This process was repeated an additional 4 times. Viral RNA was purified from supernatants of passages 1 through 5 using the ZR Viral RNA Kit (Zymo Research). The proportions of wild type and Y275H virus in each viral RNA sample was determined by digital droplet RT-PCR with the QX100 Digital Droplet PCR System (BioRad) using the One-Step RT-ddPCR Kit for Probes (BioRad) and a custom TaqMan SNP detection assay from Life Technologies (Part Number 4332077). This assay included non-specific poliovirus primers (5’- CGGAGACAGAGTTGACTACATCGA-3’ and 5’- CCGCCCTTGACACAGTATGTTTTAT-3’) and specific probes targeting wild type and Y275H virus sequences (WT: 5’-VIC- CACACCACCTGTACAAGA-NFQ-3’ and Y275H: 5’-FAM- CACCACCTGCACAAGA-NFQ-3’).

For sanger sequencing-based detection, HeLa S3 cells were coinfected with wild type and either Y275H or D79H virus at m.o.i. 0.05 each for 8 hours. Cells were frozen and thawed 3 times and cleared at 3500 rpm for 5 minutes. To maintain a low m.o.i., 1% of the viral supernatants were transferred to fresh monolayers of HeLa S3 cells and incubated for 8 hours. This process was repeated an additional 6 times for 8 passages total. Viral RNA was purified from passage 1 and 8 supernatants by TRIzol (Invitrogen) extraction and ethanol precipitation. RNA was reverse transcribed using Thermoscript Reverse Transcriptase (Invitrogen) and the 3D^pol^ region was amplified using Phusion polymerase (New England BioLabs). The amplicons were sanger sequenced to determine the proportions of wild type and mutant viruses in each population.

### Ribavirin resistance assay

HeLa S3 cells were pretreated with 0, 200, 400, 600, 800 or 1000 µM ribavirin for 4 hours then infected in triplicate with wild type, Y275H, G64S or H273R at m.o.i. 0.1 for 30 minutes. Infected cells were washed 2 times with PBS and covered with growth medium containing the same concentrations of ribavirin as used for pretreatment. Twenty-four hours post infection, cells were frozen and thawed 3 times and cleared for 5 minutes at 21000 g. Supernatants were titered by plaque assay.

### CRE-REP assay

*In vitro* transcribed RNA from either pT7Rep3L and pT7/SL3 or pRLucWT and pT7/SL3 were cotransfected (250 ng each per well in a 12-well plate) into 80-90% confluent L929 cells using Lipofectamine 2000 following the manufacturer’s instructions. Plates were harvested 48 hours post transfection and cleared briefly at 1200 rpm. Recombinant virus in the supernatant was quantified by plaque assay in HeLa cells.

### Expression and purification and WT and Y275H PV 3Dpol

The Y275H mutation was introduced into the bacterial expression plasmid (52, 53) for PV 3Dpol by using quickchange site directed mutagenesis. Expression and purification of WT and Y375H PV 3Dpol was performed essentially as described previously (52, 53).

### Template-switching assay

Elongation complexes were assembled by incubating 1 μM active-site titrated WT or Y275H PV 3Dpol with 20 μM sym/sub-U RNA primer-template (10 μM duplex) (26) and 500 μM ATP for 3 min. Template-switching reactions were initiated by addition of 60 μM RNA acceptor template (5’-GCAAGCAUGCAUGG-3’) and 500 μM CTP, GTP and UTP and then quenched at various times by addition of 50 mM EDTA. All reactions were performed at 30 °C in 50 mM HEPES, pH 7.5, 10 mM 2-mercaptoethanol, 60 μM ZnCl_2_, and 5 mM MgCl_2_. Products were analyzed by denaturing PAGE. Gels were visualized by using a PhosphorImager and quantified by using ImageQuant software (GE Healthcare).

### Infection of susceptible mice

cPVR mice (33) were infected intracranially (i.c.), intramuscularly (i.m.) or intravenously (i.v.) with either wild type or Y275H virus. I.c. infections were performed using 2-week-old mice with 5x104 plaque forming units (p.f.u) per mouse (11 mice per virus strain, 5 µl per mouse). I.m. infections were performed using 8 to10-week-old mice with 10^6^ p.f.u. per mouse (25 mice per virus strain, 50 µl per hind limb). I.v. infections were performed using 6-week-old mice with 3x10^8^ p.f.u. per mouse (15 mice per virus strain, 100 µl per mouse injected into the tail vein). Mice were monitored daily for signs of paralysis. Mice were euthanized upon appearance of dual hind limb paralysis, a sign of imminent death, and death was recorded for the following day.

## Acknowledgements

We thank Ibrahim Moustafa for contributing Figure 7. This work was supported by NIH (R01, AI36178, AI40085, P01 AI091575) and the University of California (CCADD), and DoD-DARPA Prophecy to RA. AA was supported by NSF GRF. JJA and CEC were supported by grant AI045818 from NIAID.

---

**Figure S1.**
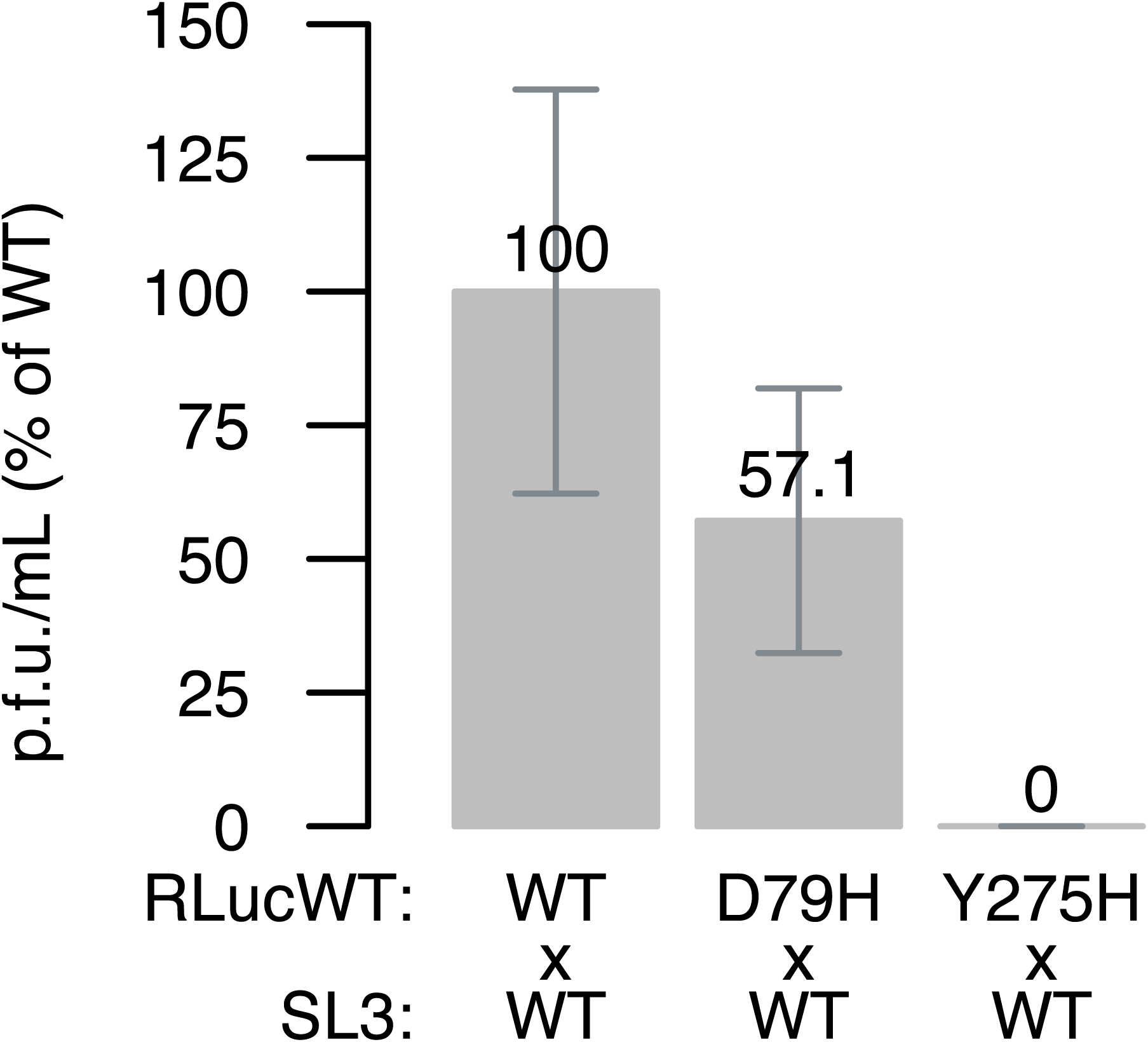
Y275H exhibits a stronger defect in recombination than the previously characterized D79H mutation (21, 22). An intertypic CRE-REP assay was used to measure relative titers of viable, recombinant progeny after a single round of replication for both Y275H and D79H. Titers are normalized to RLucWT (poliovirus type 1 Mahoney) and SL3 (poliovirus type 3 Leon) RNAs containing wild type viral polymerase. Error bars represent standard deviations of three replicates.

**Figure S2.**
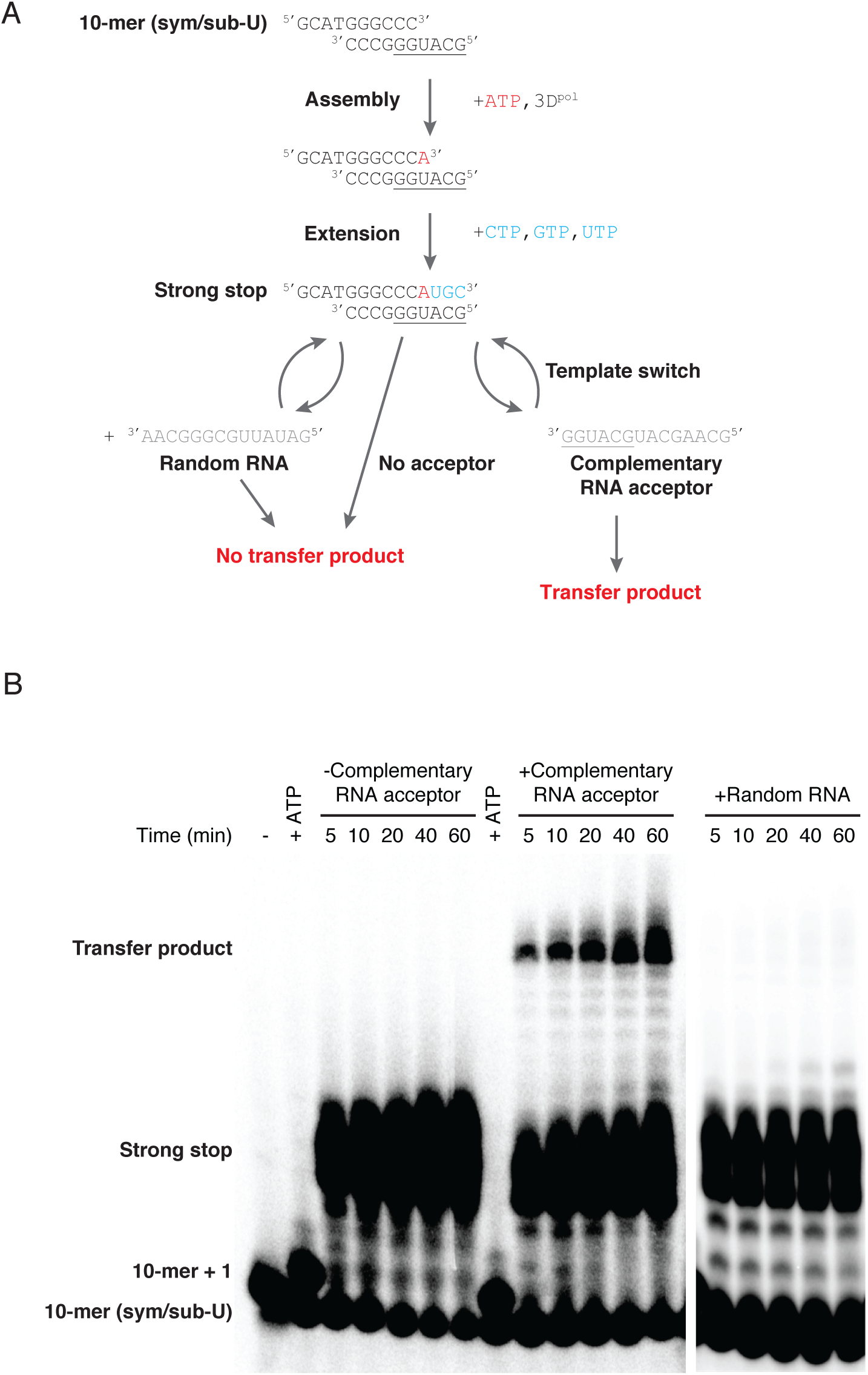
Homologous template RNA is required for template-switching *in vitro*. (A) Scheme for *in vitro* template-switching assay with random RNA acceptor, no acceptor or complementary RNA acceptor. No transfer product should be produced in the absence of complementary acceptor RNA. (B) Elongation complexes formed with 1 μM sym/sub-U, 5 μM 3Dpol and 500 μM ATP are elongated by the addition of 500 μM each of CTP, GTP and UTP in the presence or absence of RNA acceptor partially homologous to the sym/sub-U template strand or presence of a random RNA acceptor. Accumulation of high molecular weight RNA (transfer product) in the presence of acceptor RNA indicates the occurrence of template switching.

**Figure S3.**
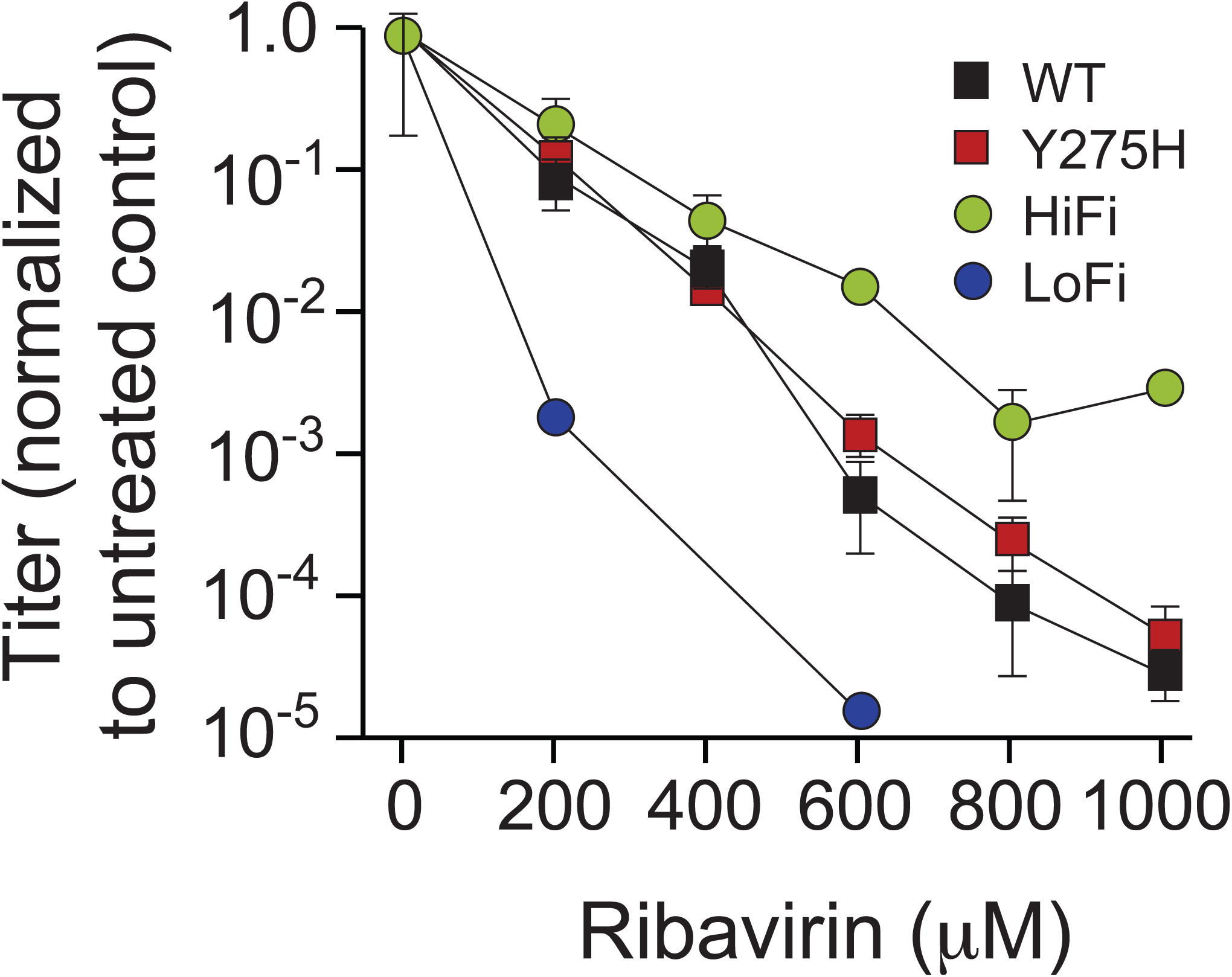
Y275H does not significantly alter replication fidelity. Wild type (square black), Y275H (square red), G64S (HiFi, circle black) and H273R (LoFi, circle red) viruses are used to infect cells treated with varying concentrations of ribavirin. Viral titers are normalized to the 0 μM ribavirin treatment for each group. Error bars represent standard deviations of three replicates. No virus was detected for the H273R virus at 400, 800 and 1000 μM ribavirin.

**Figure S4.**
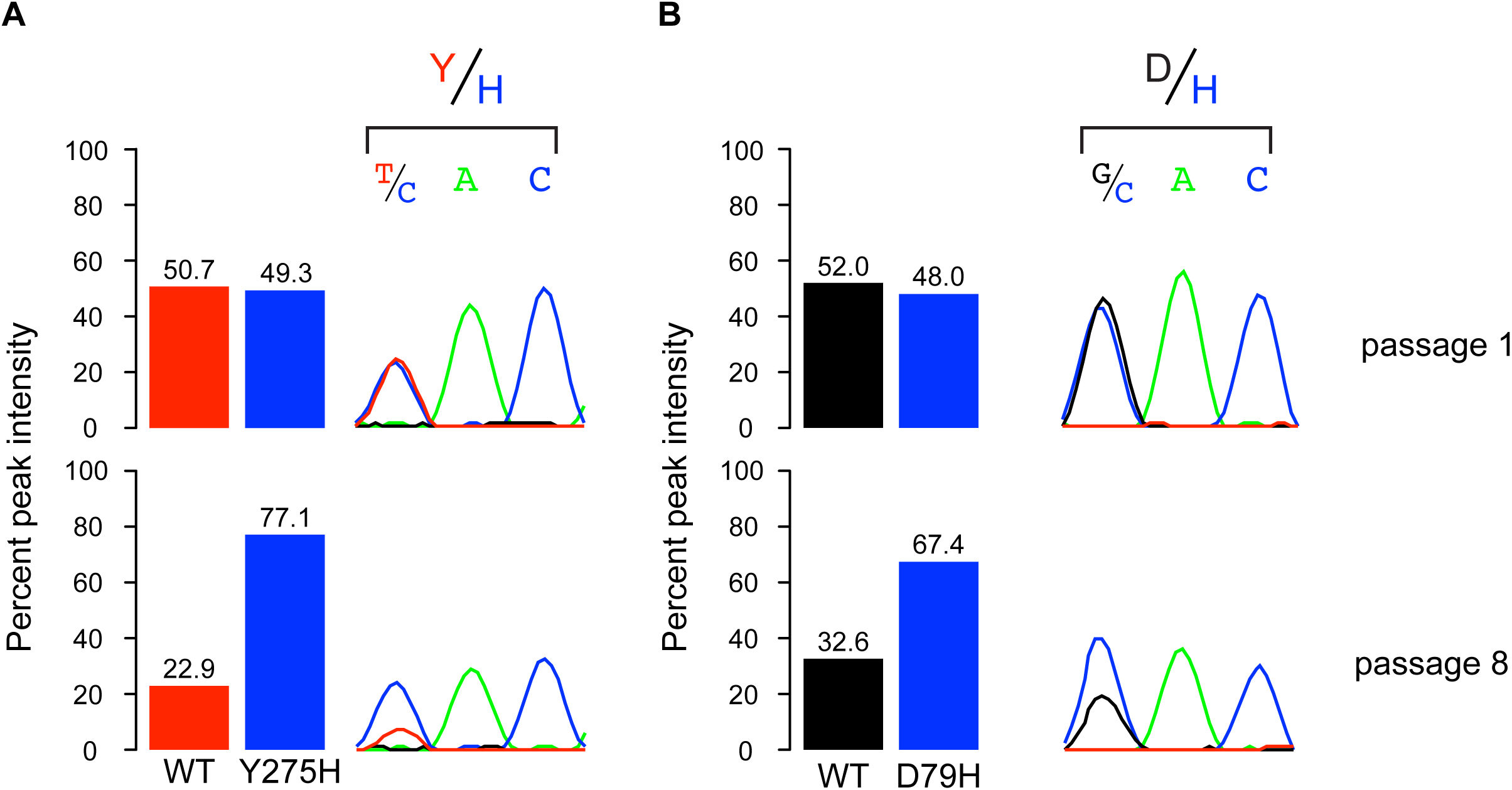
Y275H and D79H both confer a fitness advantage over wild type virus in HeLa S3 cells. (A) Y275H and (B) D79H were each initially mixed 1:1 with wild type virus and passaged 8 times at low m.o.i. The peak intensities of sanger sequenced RT-PCR amplicons of passaged populations was used to approximate the relative abundance of wild type and mutant viruses. Passage 1 peak heights for wild type and each mutation are equivalent. After 8 passages, both mutants outcompete wild type virus. Chromatograms are shown for the Y275 and D79 codons. Chromatogram and barplot colors correspond to nucleotide bases A (green), C (blue), G (green) and T (red).

